# Methanol Crude Extract of *Litsea Monopetala* Leaves combats oxidative stress, clot formation, inflammation and stool frequency in Animal Model

**DOI:** 10.1101/2024.10.31.621363

**Authors:** Zubair Khalid Labu, Samira Karim, Sarder Arifuzzaman

## Abstract

*Litsea monopetala* (LM) leaves are used in traditional medicine systems in the South Asian region for treating ailments such as digestive issues, respiratory problems, and skin disorders. In this study, we first investigated the possible antioxidant, thrombolytic characteristics, analgesic, and antidiarrheal activity of the methanolic crude extract of LM leaves. To assess the antioxidant, thrombolytic, analgesic, and antidiarrheal activity of the crude extract we used DPPH free radical scavenging, clot lysis, acetic acid, and castor oil-induced animal model, respectively. Before applying the extract to the animal model, phytochemical screening was performed to estimate the bioactive compounds (e.g.- phenol, flavonoid, alkaloids, tannins, and others) present. The extract exhibited DPPH free radical scavenging features with an IC_50_ of 8.99 µg/mL compared to ascorbic acid, an IC_50_ of 13.38 µg/ml. At 500 mg/kg dose, the extract produced a significant decrease (67.05%) in the frequency of writhings while diclofenac sodium decreased by 74.15%. The extract also significantly (*P<0.01*) decreased the frequency of diarrhea when compared to the standard drug of loperamide (3mg/kg). Finally, we assessed the clot lysis function as representative of the thrombolytic activity of extracts in comparison to streptokinase and demonstrated clot lysis action (74.15% decrease). From a therapeutic implication, we suggest that crude extract of LM’s leaves may contribute to the alternative or additive strategy to modulate conditions such as oxidative stress, clot formation, inflammation and diarrhea.

## Introduction

*Litsea* genus pant *Litsea monopetala* (LM) is a well-known in tropical and subtropical regions including Bangladesh (Fig.1). It is an evergreen tree growing from 5‒20 metres tall. The bole is usually straight or crooked and is up to 60cm in diameter. Branchlets are densely ferruginous pubescent. Leaf blades are broadly oval or obovate to ovate-oblong, thickly ferruginous pubescent abaxially, along midrib ferruginous pubescent adaxially when young, pinninerved, with 8–12 pairs of lateral veins, a base that is rounded or acute, and an apex that is obtuse or rounded. The leaves alternate, and the petiole is 1-3 cm, densely hairy like branchlets [1]. The plants in the genus *Litsea* contain a multitude of biologically active substances, including butanolides found in the leaves of *Litsea acutivena* [2], flavonoids found in the leaves of *Litsea coreana* and *Litsea japonica* [3], sesquiterpenese found in the leaves and twigs of *Litsea verticillata* [4], and essential oils found in the leaves, LM fruits, flowers, and bark of LM, and the fruits of *Litsea glutinosa* [5–7].

**Fig. 1.**
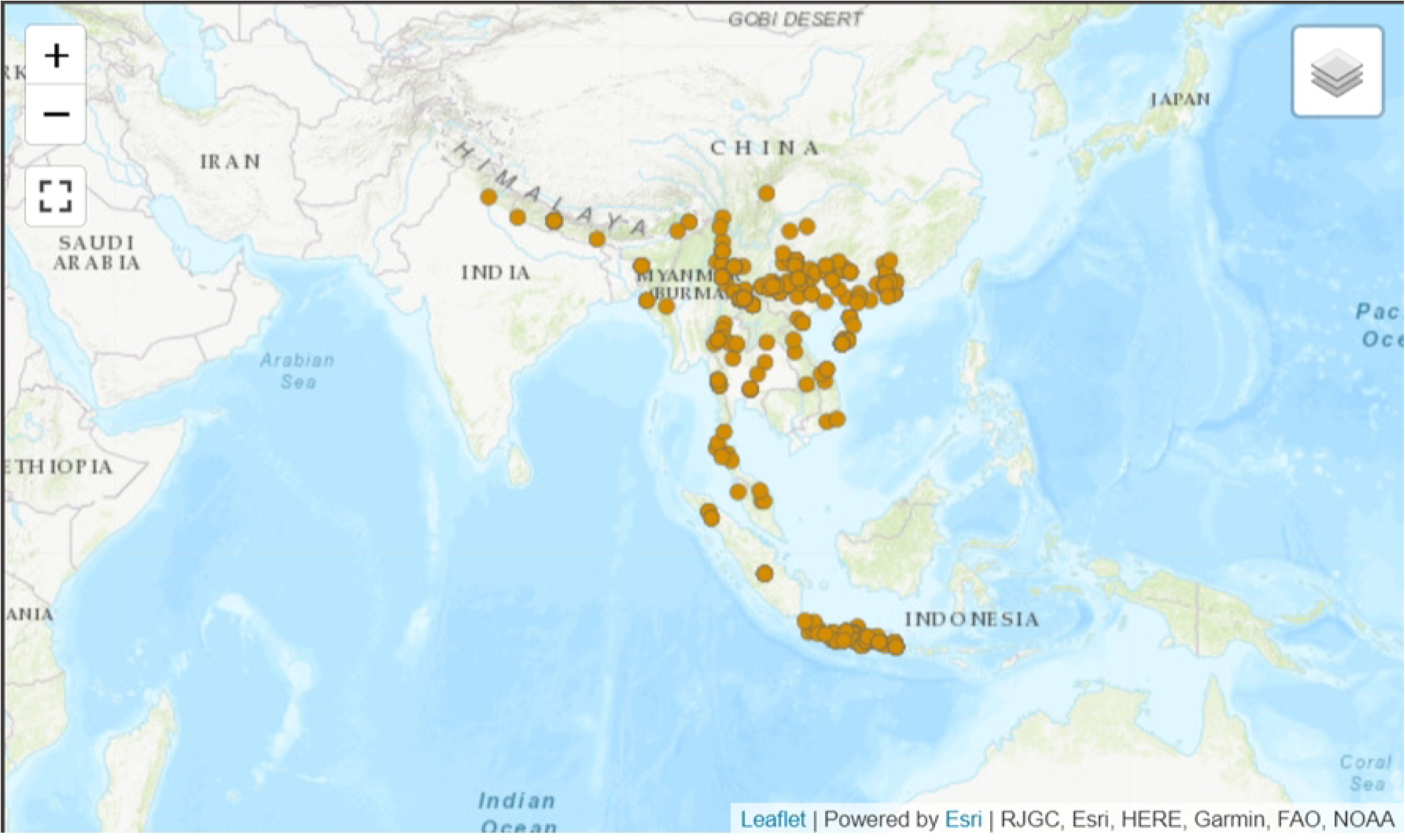
Distribution of *litsae monopetala* plant in the planet.

There are reports that the LM seed contains around 30% oil which is used for industrial purposes [8]. Extract of the LM leaves is used as an ingredient in commercial cosmetic preparations as a skin conditioner [9]. The leaves are also used as a topical medicine for the treatment of arthritis[10]. The bark of this species is mildly astringent and powdered bark and roots are used externally against bruises and pains. Leaves are used for growing muga silkworm and as cattle fodder. External application of leaf, bark, and root powder is used to treat bruising and aches. LM herbs are also used to treat animal fractures[11, 12]. There have been reports of antioxidant [12–16], antihyperglycemic [17], and analgesic [16] properties in leaves.

Recently, in an *in-vitro* study, Lamichhane G *et al.* suggested that with the selected five plants LM leaves also possess remarkable antioxidant, antibacterial, anti-adipogenic, and anti-inflammatory activities [12]. The biological activity of crude extracts of the genus *Litsea*, and the phytochemicals isolated from these extracts, has also been reported. Arfan *et al* evaluated the use of phenolic compounds isolated from the bark extract of *LM* for their antioxidant activity [13]. *LM* (Roxb.) Pers. is used in cattle diarrhea, stomach ache, gastric ulcers, and bone fractures [16, 18]. To date, no study conducted in *in-vivo* experiments to find possible therapeutic benefits for common health problems. Thus, we have attempted to evaluate the antioxidant, analgesic, antidiarrheal and thrombolytic properties of LM leaf methanolic extract in animal models.

## Materials and Methods

### Ethical approval

The animal experiment was done by following instructions followed by the ICDDRB that were approved by the Institutional Animal Ethical Committee. World of University Ethical Committee approved the experiment.

### Experimental animal

For this experiment, Swiss albino mice, aged 4-5 weeks, with a mean weight of 20-25 g, were used in this study. All the animals were bought from Jahangirnagar University (JU), in Bangladesh’s Animal Research Lab. They were given JU-formulated food and water and kept in a typical setting for a week at the World University of Bangladesh research facility to help them acclimate.

### Chemicals

We purchased quercetin, ferric chloride, gallic acid, trichloroacetic acid and DPPH (1,1-diphenyl, 2-picryl hydrazyl) from Sigma Chemical Co (USA). Potassium ferricyanide was purchased from May & Backer, Dagenham, UK, and ascorbic acid was acquired from SD Fine Chem. Ltd. in India. We bought potassium ferricyanide, sodium carbonate, methanol, and Folin-Ciocalteu reagent from Merck in Germany. Lyophilized streptokinase (1,000,000 IU) was purchased from Square Pharmaceuticals Ltd., Dhaka, Bangladesh. All the chemicals used in this experiment were analytical grade.

### Collection and identification of plant sample

The fresh plant was collected for Tangail district village, Bangladesh. Freshly collected branches and leaves were taken to the Bangladesh National Herbarium in Mirpur, Dhaka for authentication. The voucher specimen was added to the Bangladesh National Herbarium and given the accession number 45413.

### Drying and grinding of plant materials

The collected leaves were cleaned under clean running water to remove any remaining dirt. The samples were first allowed to dry at room temperature under the shade for a week, then dried at 50 to 60°C in a mechanical dryer to achieve complete drying. The dried leaves were mechanically ground into a coarse powder. The powdered sample was stored in a sealed airtight container in a cool, dry, and dark place till further use.

### Extract Preparation

Four hundred gram (400 g) of leaf powder was weighed and dissolved in 1500 ml methanol, and extracted using a Soxhlet extraction apparatus for 72 h. The methanol solvent was evaporated from the extraction by using a rotary evaporator under reduced pressure to obtain methanol crude extract. A greenish-black gooey and sticky concentrated methanol crude extract was used for phytochemical screening and pharmacological activity evaluation. The extract was stored at 40°C in a Pharmaceutical standard refrigerator until further use. Six grams of crude extract dissolved and extracted with aqueous methanol (MSF), petroleum ether (PSF), Ethyl acetate (ESF), carbon tetrachloride (CTF), and chloroform (CSF). The fractionated amounts were PSF(1.5g), ESF(1.5g), CTF(1.4g), CSF(0.5g), and residual MSF(0.11g), respectively. All crude extracts were filtered separately through Whatman No. 41 filter paper to remove particles. The particle-free crude extract was evaporated completely by using rotary evaporator under reduced pressure to obtain dry crude extracts. The residue left in the separatory funnel was re-extracted twice following the same procedure and filtered. The combined extracts were concentrated and dried by using a rotary evaporator under reduced pressure.

### Phytochemical screenings

All crude extracts were subjected to confirmatory qualitative phytochemical screening with a slight modification of Sofowara’s standard protocol for phytochemicals in the crude extracts [19]. The main classes of compounds obtained were various organic soluble fractions, including alkaloids, flavonoids, steroids, phenols, resins, glycosides, and saponins, as well.

### Estimation of total phenolic content (TPC)

By utilizing the Folin-Ciocalteu [20] reagent, the TPC was determined. Two milligrams of crude extract were liquefied with 2 mL of distilled water. In the test tube, 2 mL of diluted Folin-Ciocalteu (previously diluted 10-fold with deionized water) was mixed with 0.5 mL of extract (1 mg/mL). The mixture was left to stand at 22 ± 2 °C for five minutes. Then 2.5 milliliters of 7.5% Na2CO3 was added to the test tube. To allow for color development, the mixture was gently mixed for 20 minutes without allowing any significant jerking. The UV-Vis spectrophotometer (Model: UV-1700 series) method was used to measure the color change’s intensity at 760 nm. The compound’s TPC was represented in the absorbance value. The final concentration of 0.1 mg/mL was used to assess the extract and standard sample. The GA equivalent, or GAE (Standard curve equation: y = 0.0085x +0.1125, R² = 0.9985), was used to express TPCs as mg of GA/g of dry extract, Fig. 2.

**Fig. 2.**
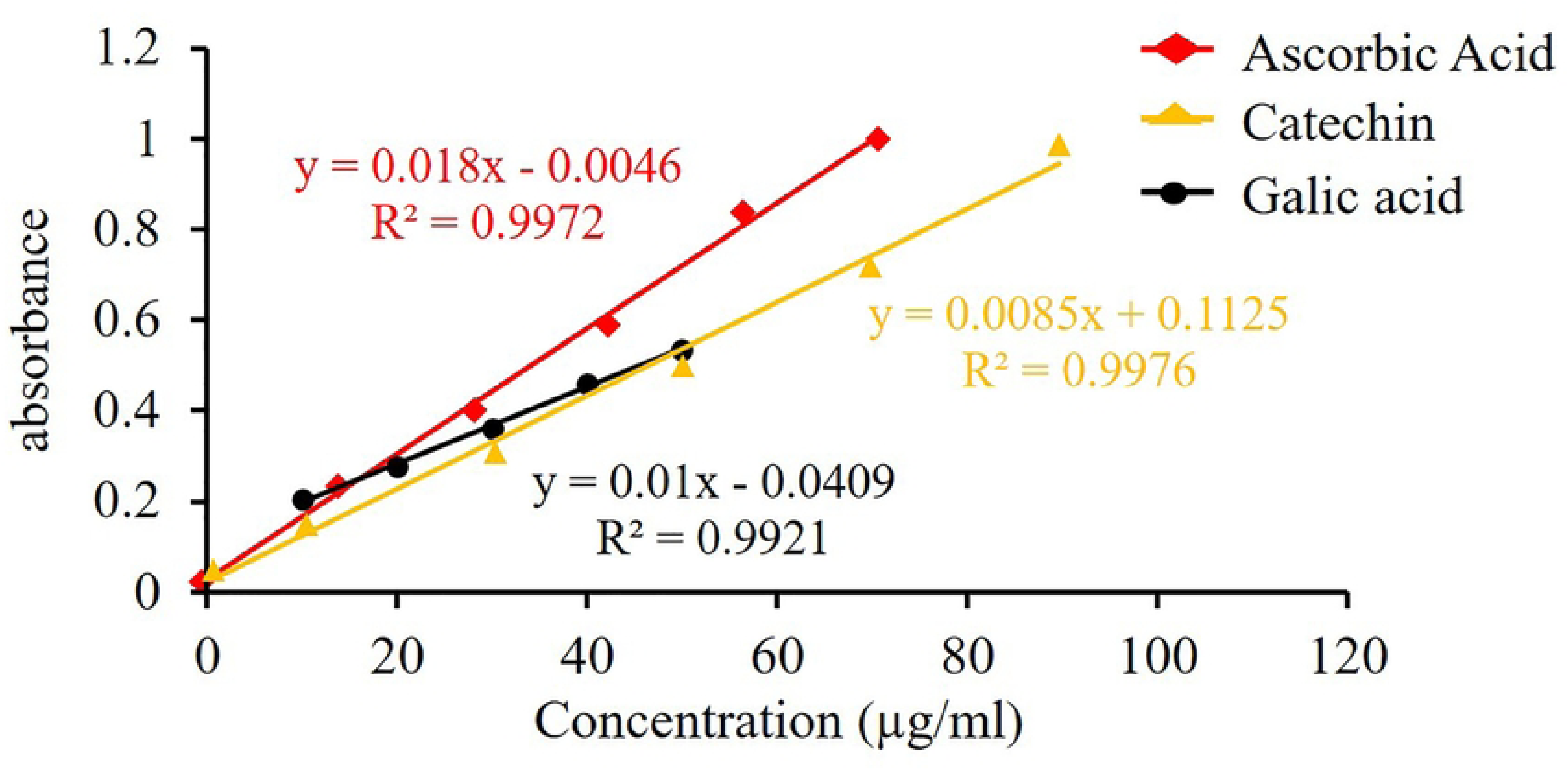
Standard curve of Gallic acid, Catechin and Ascorbic Acid.

### Determination of total flavonoid content (TFC)

We followed the aluminum chloride colorimetric method, the total flavonoid concentration of the crude extracts was determined [21]. In brief, 0.1 mL of 10% AlCl_3_ was added with 1.5 mL of crude extract, and then 0.1 mL of 1 M Na-acetate was subsequently added to the reaction mixture. The mixture was left to stand for half an hour. After that, 1 mL of a 1 mol/L NaOH solution was added, and double-distilled water was used to bring the mixture’s final volume to 5 mL. After the mixture was let to stand for 15 minutes, the absorbance at 415 nm was determined. The calibration curve was used to determine the total flavonoid content, which was then reported as mg of quercetin equivalent per g of dry weight. Total flavonoid content was calculated from the following calibration curve: Y = 0.01 X + 0.0409, R2 = 0.9921, where Y is the absorbance of crude extract and X is the quercetin equivalent, Fig. 3.

**Fig. 3.**
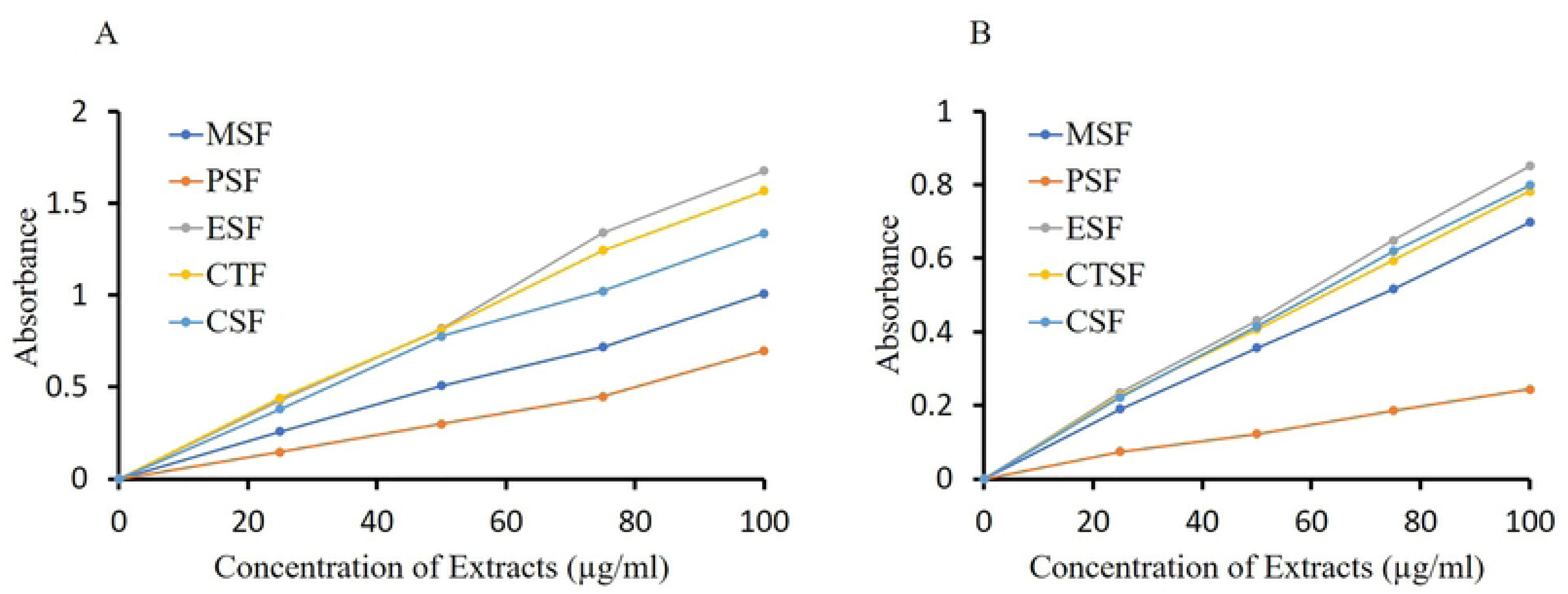
Estimation of (A) Total phenolic contents; (B) Total flavonoid contents among various fractions of *LM* leaves.

### Acute toxicity test

To test acute toxicity, the methanol crude extract was administered to mice at doses 200, 400, 800, 1600, and 3200mg/kg p.o. We used six mice in each group. Prior administration to mice, animals were kept fast for 16h. All animals were allowed free access to food and water followed by observed for 48h for symptoms of acute toxicity. The number of deaths within this period was recorded [22].

### Antioxidant activity assay

We followed the technique outlined by Williams et al. [20], and the free radical scavenging activities (antioxidant capacity) of the extracts on the stable radical DPPH were calculated. The ability of DPPH to scavenge free radicals was demonstrated by the spectrophotometrically magnified change in colour from purple to yellow in methanol. 3.0 mL of DPPH methanol solution (20 µg/mL) was combined with 2.0 mL of MSF and its fractions of extracts at varying concentrations. By using a UV spectrophotometer to compare the extract’s antioxidant capability to ascorbic acid (AA), the extract was shown to be capable of bleaching a purple-colored methanol solution containing DPPH radicals. Positive control was employed with AA. The mother solution with a concentration of 1000µg/mL was created by dissolving a calculated quantity of AA in methanol. The mother solution was serially diluted to get differential concentrations ranging from 500.0 to 0.977 µg/mL. The mother solution (1000 µg/mL) was obtained by dissolving the measured quantity of various extracts in methanol. Differential concentration was obtained by dilution of the mother solution in steps. 20 µg/mL of DPPH solution was obtained by weighing and dissolving 20 mg of DPPH powder in methanol. The prepared solution was stored in the light-proof box together with the amber reagent bottle. 3.0 mL of DPPH methanol solution (20 µg/mL) was combined with 2.0 mL of a methanol solution of the sample (extractives/control) at varying concentrations (500 µg/mL to 0.977 µg/mL). Using methanol as a reference, the absorbance was measured at 517 nm using a UV spectrophotometer following a 30-minute reaction at room temperature in a dark environment. The following formula was used to get the percent inhibition of free radical DPPH inhibition:

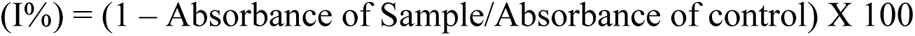

Similar procedure was repeated for the standard ascorbic acid instead of sample solution (plant extract) to obtain the percentage inhibition. The graph showing the proportion of inhibition against extract concentration was used to determine the extract concentration that would provide 50% inhibition (IC_50_).

### Analgesic activity assay

To test the analgesic activity, experimental animals are randomly divided in 3 groups (n=6). Each group received different treatment: the extract, the standard (diclofenac sodium), and the control. Diclofenac (standard) and test samples (plant extract) are administered orally using a feeding needle. All the experimental animals received an intraperitoneal injection of 0.7% acetic acid solution (15 ml/kg), a writhing-inducing chemical, to guarantee adequate absorption of the drugs provided after a 30-minute interval. Acetic acid activates the nerve and is used to simulate writhing by releasing endogenous chemicals that cause algia [36]. Following a 30-minute injection (IP) of acetic acid, their body contractions are observed. Then the test group received crude extract at a dose of 500 mg/kg BW and the standard group received diclofenac at a dose of 25 mg/kg BW, while the control received a diluted tween-80 solution. After five minutes, the writhing number (squirms) was meticulously counted for 15 minutes. All test samples are adjusted by diluting with distilled water to 5.0 ml volume.

### Antidiarrheal activity assay

To assess the antidiarrheal activity of the crude extract we adopted a previously published method by Shoba and Thomas et. al. [23]. Experimental animals are grouped, each consisting of six mice. The group under control: received vehicle (1% Tween 80 in water) orally at a dose of 10 ml/kg body weight; the positive control group received loperamide orally at a dosage of 3 mg/kg body weight. Test group 1 received an oral dosage of 200 mg/kg body weight of an ethanolic extract; Test group 2 received an oral dose of 400 mg/kg body weight of an ethanolic extract. Oral administration of 0.5 mL castor oil was used to produce diarrhoea in each mouse 30 mins following the aforementioned treatment. Every mouse was housed in a separate cage, and white blotting paper was used to line the floor. The experiment was carried out for four hours, changing the blotting sheets every hour. The total amount of diarrheal faeces (watery, unformed stool) that occurred throughout the observation period was noted. After an hour, the old absorbent papers were swapped out for new ones.

The control group’s total amount of diarrheal faeces was considered to be 100%. The defecation inhibition percentage was calculated using the following formula.

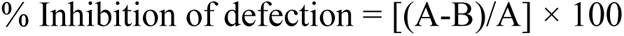

A= Mean number of defecations by castor oil; B= Mean number of defecations by extract/ loperamide

### Castor oil-induced Enteropooling assay

We assess the intraluminal fluid buildup following the Robert et al. (1976) method. Experimental animals are randomly divided into four groups and six animals in each group. The Group 1 and 2 received prostaglandins (200 micrograms/kg; ip) and distilled water (Oral), respectively. The group 3 and 4 received crude extracts orally at doses of 300 and 600 mg/kg, respectively, one hour before the castor oil was administered orally. The mice were put to death and given ether anaesthesia two hours later. The small intestine was removed and weighed after the margins were secured with thread. Squeezing the intestinal material into a graduated tube allowed for the measurement of its volume. The difference between the full and empty intestines was computed after the intestine was reweighed [24].

### *In-vitro* thrombolytic activity test

A method published by Prasad et al. [25] was followed to assess the thrombolytic activity of the crude extract. The protocol was approved by the ethical committee of the World University of Bangladesh, Dhaka, Bangladesh. Lyophilized Streptokinase powder was purchased from Beacon Pharmaceuticals Ltd, Dhaka, Bangladesh. Five milliliters of sterile distilled water was added to the vial to reconstitute the streptokinase and mixed thoroughly. The reconstituted sample was stored as per the labeled instructions. Venous blood was drawn from the healthy human volunteers (*n*=10) and transferred (0.5 ml/tube) to the previously weighed sterile microcentrifuge tube to form the clot. Before withdrawing the blood sample, a written agreement was taken from the volunteers. The samples are incubated at 37 °C for 45 min. After the clot formation, serum was completely removed without disturbing the clot and each tube having clot was again weighed to determine the clot weight. A volume of 100 µl of methanol extract (10 mg/ml) was added to each microcentrifuge tube containing pre-weighed clot. As a positive control, we added 100 µl (30000 IU) of streptokinase, and as a negative control 100 µl of distilled water to the control tube. All the tubes were then incubated for 90 minutes at 37 °C and observed the clot lysis. After incubation, the fluid released was removed and tubes were again weighed to observe the difference in weight of clot lysis. The difference obtained in weight taken before and after clot lysis was expressed as a percentage of clot lysis. All the experiments are performed in at least three replicates.

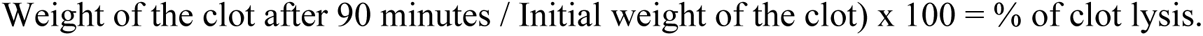

### Statistical analysis

Results were expressed as mean ± SEM. To determine the statistical significance one way ANOVA followed by Dunnett’s multiple comparisons was performed.

## Results and discussion

### Identification and Confirmation Crude Phytochemicals present in the LM extract

Several phytochemicals, including tannins, phenols, steroids, flavonoids, alkaloids, saponins, and other phytocompounds, were confirmed in the extract through qualitative tests (Table 1). Among the tested phytochemicals our results reveal tannins as the major phytocompound present in the crude extract as determined by the lead acetate test. Qualitative tests also demonstrate a moderate presence of carbohydrates, alkaloids, and flavonoids in the extract as determined by the molisch’s test, mayer’s test, and shinoda’s test, respectively. While reducing sugars, steroids, and phenolic content were present in insignificant quantities. Interestingly, saponins was absent in the extract.

**Table 1.** Phytochemical screening of methanolic extract of LM leaves.

| Phytochemicals | Test Method | Remarks | Description |
| --- | --- | --- | --- |
| Carbohydrates | Molisch's Test | ++ | Moderately present |
| Reducing sugars | Fehling solution test | + | Mildly present |
| Tannins | Lead acetate test | +++ | Largely present |
| Alkaloids | Mayer's test | ++ | Moderately present |
| Flavonoids | Shinoda test | ++ | Moderately present |
| Saponins | Frothing test | - | Not present |
| Steroids | Liebermann-Burchards test | + | Mildly present |
| Phenol | Ferric chloride test | + | Mildly present |

### Estimation of total phenolic contents (TPC) in Crude extract

Using gallic acid (GAE) as the standard, we observed a straight-line Y = 0.01 X + 0.0409, R2≥0.9921 graph of the TPC in the crude methanol extract (Fig. 3). In particular, we tested the TPC in MSF, PSF, CTF, CSF, and ESF of leaves of LM crude methanol extract. The average TPC found were 165.02, 65.15, 219.18, 185.73, and 301.67 mg/g of extract, respectively (Fig. 3C, Table 2). We observed a lower quantity of TPC in PSF (65.15 ± 0.27 mg/g) while in ESF (301.67 ± 0.40 mg/g) TPC obtained was maximum, followed by CTF and CSF. In comparison to MSF and PSF, the phenolic contents of ESF were significantly higher (p < 0.05). This finding implies that all of the soluble fractions of LM leaves offer phenol content at varied degrees. The results of this study show that LM flowers might have a good source of phenolic compound with possible health advantages.

**Table 2.** Total phenolic, total flavonoid contents of crude extract of LM (TPC, Total phenolic content; TFC, total flavonoid content; MSF, Crude methanol extract; PSF, Petroleum ether fraction; ESF, Ethyl acetate fraction; CTF, Carbon tetrachloride fraction; CSF, Chloroform fraction).

| Sample | TPC (mg/ml)<br>Mean±SEM | TFC (mg/ml)<br>Mean±SEM |
| --- | --- | --- |
| MSF | 165.02 ±0.57 | 686.40 ± 0.52 |
| PSF | 65.15 ± 0.55 | 165.02 ± 0.45 |
| ESF | 301.67 ±0.47 | 635.41 ± 0.54 |
| CTS | 219.18 ± 0.37 | 699.30 ± 0.61 |
| CSF | 185.73 ± 0.15 | 702.40 ± 0.35 |

### Estimation of total flavonoid content (TFC)

Flavonoids are well known for their antioxidant properties, thus attracting considerable interest in medicinal plant search research. We used the complexometric technique to quantify total flavonoid concentration in the extract (Fig. 3B). Like TPC, we observed a straight-line (Y = 0.01 X + 0.0409, R2 ≥0.9921) graph for the TFC concentration in the extract. We used catechin (with a concentration range of 10.0 μg/mL to 160.0 μg/mL) as the standard, and the flavonoid contents were evaluated in terms of catechin equivalent (CE) (Fig 3). The TFC was measured in the presence of other organic solvents, including PSF, CTF, CSF, and ESF. The average TFC found were 686.30, 165.01, 635.10, 699.30, and 702.40 mg/g of dry extract, respectively (Table 2). The maximum amount of TFC was found in ESF (702.30 ± 0.35 mg/g) and the lowest amount in PSF (165.35 ± 0.45 mg/g), followed by CSF, CTF, and MSF. When compared to MSF and PSF, the flavonoid contents of ESF were considerably higher (p < 0.05). The study’s confirmation of noteworthy flavonoid concentrations in all soluble fractionates suggests that LM leaves are a rich source of flavonoids, which makes them useful for medicinal purposes.

### Effect of LM crude Extract on total antioxidant capacity (TAC)

We evaluated the TAC of the crude extract in comparison to the ascorbic acid (AA). A complexometric technique involving the reduction of molybdenum (V) to molybdenum (III) was used to measure the TAC (Fig. 4). Based on a known concentration of AA, a straight-line graph was obtained (Fig. 4). The TAC found in MSF, PSF, CTF, CSF, and ESF were 329.61 ± 0.19, 132.51 ± 0.51, 318.04 ± 0.71, 399.76 ± 0.19, and 400.90 ± 0.59 mg/g of dry extract, respectively (Table 3). Except PSF, CSF, CTF, MSF and ESF showed the greater TAC (>400.90 mg/g of dry extract) that were almost close to AA. In both ESF and CSF, TAC was considerably greater (p < 0.05). The TPC and TFC are connected to the TAC of each fraction. TPC and TFC exhibited a significant connection with total antioxidant capacity (r2 = 0.946, 0.965; p < 0.05) according to Pearson correlation analysis (Supp. Table S1). By donating a proton, plant polyphenols and flavonoids balance singlet or triplet oxygen and may shield the body from oxidative stress [26].

**Fig. 4.**
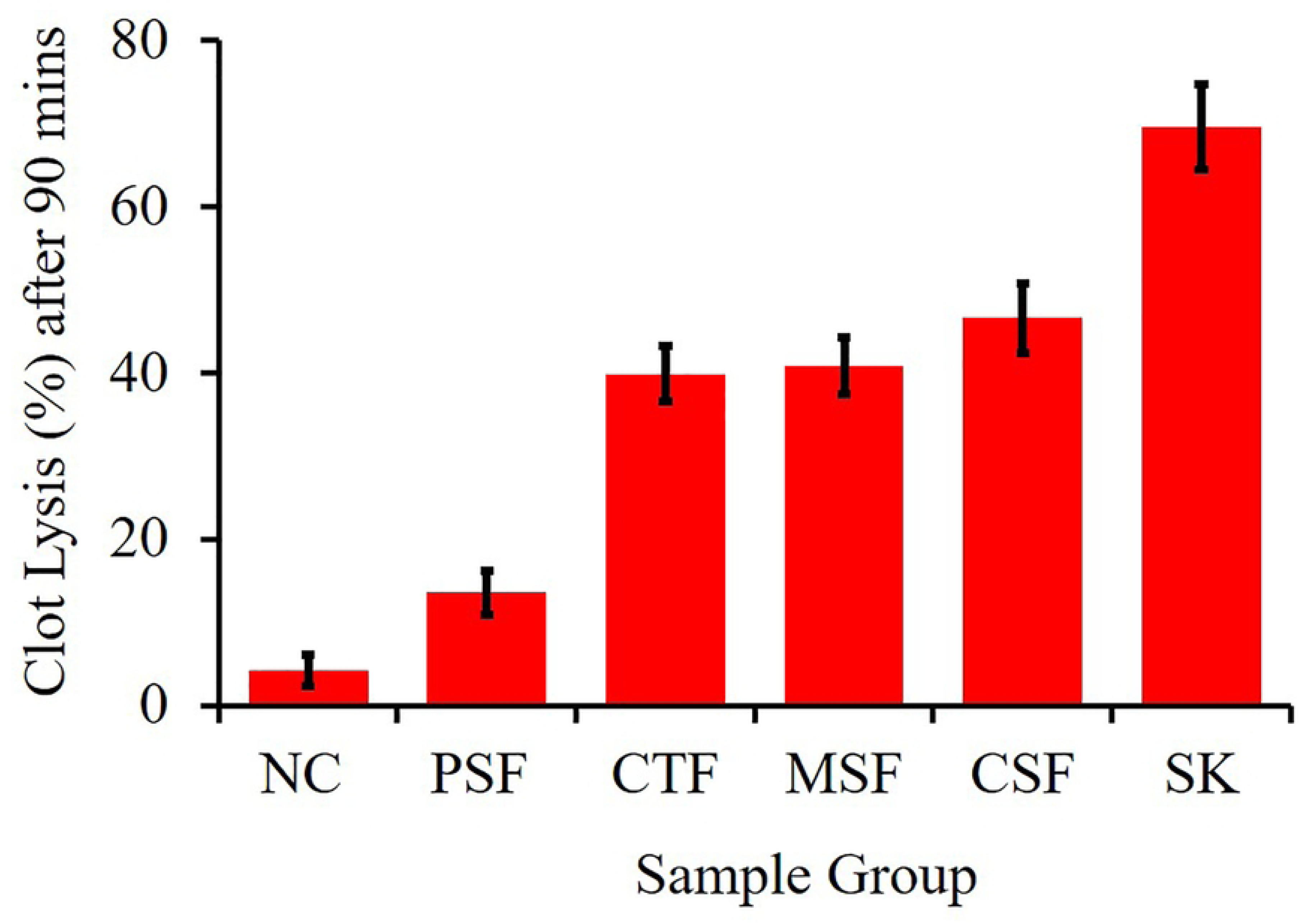
In vitro analysis of antioxidant activity among various fractions of leaves of *litsae monopetala*. (A) Total antioxidant capacity analysis; (B) DPPH free radical scavenging activity analysis. Ascorbic acid (AA) was used as the standard antioxidant. CSF, Chloroform fraction; CTF, Carbon tetrachloride fraction; ESF, Ethyl acetate fraction; MSF, Crude methanol extract; PSF, Petroleum ether fraction.

**Table 3.** Total antioxidant capacity and DPPH free radical neutralizing property (IC_50_) of crude extract. (The letters a, b, c, d in the superscript indicates the level of significance, as determined by Duncan’s Multiple Range Test (DMRT).

| Extract/<br>Standard | TAC<br>(mg AAE/g of extract) |  | IC<sub>50</sub> value<br>(µg/ml) |  |
| --- | --- | --- | --- | --- |
|  | Mean ± SD | *p-value* | Mean ± Sd | *p-value* |
| MSF | 329.78 ± 0.61<sup>b</sup> | 0.04 | 12.99 ± 0.21<sup>c</sup> | 0.03 |
| PSF | 132.53 ± 0.51<sup>a</sup> | 0.05 | 47.05 ± 0.46<sup>d</sup> | 0.03 |
| CTF | 318.04 ± 0.71<sup>b</sup> | 0.08 | 6.08 ± 0.21<sup>a</sup> | 0.06 |
| CSF | 399.19 ± 0.15<sup>c</sup> | 0.01 | 8.99 ± 0.21<sup>b</sup> | 0.02 |
| ESF | 400.90 ± 0.59<sup>c</sup> | 0.05 | 6.88 ± 0.411<sup>a</sup> | 0.05 |
| AA | 410.90 ± 0.39<sup>c</sup> | - | 8.99 ± 0.42<sup>b</sup> | - |

### Effect of LM crude extract on DPPH radical scavenging

We tested DPPH radical scavenging capacity of the crude extracts as the measure of the TAC (Fig. 4D). The data obtained of DPPH free radical neutralizing property (IC_50,_ %) for ascorbic acid (AA), MSF, PSF, CTF, CSF, and ESF produce a straight-line graph. For MSF, PSF, CTF, CSF, and ESF IC_50_ values were 12.99 ± 0.21, 47.05 ± 0.46, 6.08 ± 0.21, 8.99 ± 0.21, and 6.98 ± 0.41 μg/mL, respectively (Table 3). When compared to MSF, PSF, CSF, and AA, the IC_50_ values for ESF and CTF were significantly lower (p <0.05). The highest potential to scavenge free radicals caused by DPPH were CTF and ESF (IC_50_ values 6.08 μg/mL and 6.98 μg/mL, respectively), whereas AA’s IC_50_ was 8.99 ± 0.42 μg/mL. A strong association between TPC and TFC and DPPH free radical scavenging was found through correlation analysis (r^2^ = 0.9331, 0.976; p < 0.05) (Supp. Table S1). Published reports also describe the antioxidant properties of aqueous alcoholic extractives of LM’s tubers, seeds, leaves, and bark. Rich amounts of flavonoids and polyphenols in LM leaves also function as proton donors and have a strong association with the plant’s ability to scavenge free radical’s DPPH (Fig. 4).

### Effect of LM crude extract on Analgesic activity

An Acetic acid-induced writhing model was used to assess the analgesic activity of the crude extract. Acetic acid induces writhing upon the release of endogenous chemicals that excite the pain nerve and create analgesia. We used Diclofenac as a standard drug and compared the analgesia with our extract. The writhing response caused by acetic acid is significantly reduced by the LM extract at a dosage of 500 mg/kg body weight. In mice, leaves of LM induced 67.05% writhing inhibition, compared to 74.15% with conventional medicine Diclofenac at a dosage of 25 mg/kg body weight of the experimental animals (Table 4). When compared to control mice, the methanolic extract of LM demonstrated a notable analgesic effect. However, further research is needed to identify the original active ingredient that gives LM’s methanolic extracts their analgesic effects.

**Table 4:** Evaluation of the Analgesic activity using the writhing test (n=6).

| **Group** | **Writhing (Total)** | **Writhing**<br>**(Mean ± SEM)** | **Decrease**<br>**compared to**<br>**control (%)** |
| --- | --- | --- | --- |
| Control (Tween 80) | 162 | 32.5±1.85 | - |
| Diclofenac (25mg/kg) | 37 | 7.1±1.05 | 74.25 |
| Extract (500mg/kg) | 49 | 9.6±1.01 | 67.05 |

### Effect of LM crude extract on Antidiarrheal activity

We used Castor oil-induced animal models to conduct in vivo antidiarrheal experiments. Administration of crude extract to tested animals exhibited a lesser frequency of feces compared to the control group (Fig. 3). In comparison to the conventional loperamide (3 mg/kg BW) group extract showed a lower frequency of feces (Table 5). Defecation inhibition was seen in 71.1% of the extract group compared to the control group. During a 4-hour testing period, all of the extracts demonstrated antidiarrheal efficacy as evidenced by a decrease in the feces frequency in comparison to the control group. In addition to calculating the percentage of defecation inhibition, the total number of defecations throughout 4 hours was noted. The extract showed 51.09% (200 mg/kg) and 60.0% (400 mg/kg) of suppression of defecation in a dose-dependent manner. Every extract showed statistically significant inhibition of defecation frequency and volume (*p<0.05* and *p<0.01*). Previous studies on phytochemicals found that the presence of tannins, alkaloids, flavonoids, saponins, sterol, and triterpenes in medicinal plants was linked to their antidiarrheal properties [27]. Tanins cause the intestinal mucosa’s proteins to become denatured by producing protein tanuates, which increase the mucosa’s resistance to chemical modification and reduce secretion [44]. According to reports, flavonoids prevent the release of prostaglandins and autocoids, which may prevent castor oil-induced secretion and motility [28]. The study’s experimental plants are abundant in tannins, alkaloids, phenols, flavonoids, steroids, and maybe additional phytochemicals. Therefore, the presence of these beneficial phytochemicals may be the cause of the plants’ strong antidiarrheal effect [29].

**Table 5:** Effect of methanolic extract in antidiarrhoeal activity determined by feces frequency.

| **Groups** | **Dose** | **Feces Frequency at 4 hours**<br>**(Mean ± SEM)** | **Inhibition (%)** |
| --- | --- | --- | --- |
| Control | 10 ml/kg | 13.1 ± 1.1 | - |
| Positive control(loperamide) | 3 ml/kg | 2.8 ± 0.4 | 71.1% |
| Extract of LM (Test group I) | 200 ml/kg | 6.0 ± 0.6 | 51.09% |
| Leaf extract of LM (Test group II) | 400 ml/kg | 5.0 ± 0.5 | 60.0% |

**Table 6:** Effect of methanolic extract in antidiarrhoeal activity as determined by enteropooling assay. (*P<0.05, **P<0.01, Dunnett Multiple comparison test)

| Group | Volume of intestinal content<br>(ml)<br>(n=4) |  | Weight of intestinal content<br>(g)<br>(n=4) |  |
| --- | --- | --- | --- | --- |
|  | Mean + SEM | *p-value* | Mean ±SEM | *p-value* |
| Normal saline (2ml) | 1.01 ± 0.15 | - | 1.00±0.06 | - |
| Castor oil (2ml) | 2.99 ± 0.18 | - | 2.59 ± 0.5 | - |
| Extract (300mg/kg) +<br>Castor oil (2ml) | 1.18 ± 0.19 | 0.05 | 2.05±0.16 | 0.03 |
| Extract (600mg/kg) +<br>Castor oil (2ml) | 1.28 ± 0.18 | 0.01 | 2.01±0.12 | 0.01 |

We also tested antidiarrheal activity using an enteropooling assay. Oral treatment of the extract an hour before to castor oil administration substantially (P<0.05) reduced the enteropooling at 1.18 ml (300 mg/kg) and 1.28 ml (600 mg/kg). Similar to the 1.01 ml intestinal fluid volume seen in the normal group (Table 5). After receiving castor oil therapy, the weight of the intestinal content likewise dramatically increased (in normal rats, from 1.00g to 25.9.). Nonetheless, the weight of the intestinal content decreased somewhat as a result of the extract.

### Effect of crude extract on thrombolytic activity

We assessed the thrombolytic potential of the crude methanolic and another partitioning of LM. In contrast, a negative control condition of sterile distilled water investigated a small fraction of clot breaks down (3.24%). The following fractions showed a higher percentage of clot lysis in MSF (45.58%), PSF (40.00%), CTF (38.99%), and ESF (14.55%). Hence MSF exhibited a higher percentage of clot lysis (45.58%) in compare to standard streptokinase (69.52%). The interpretation of the data suggests that the LM extract had good thrombolytic activity; (p < 0.01; p < 0.05), as shown in Table 7 and Fig 5. One of the main vascular diseases that causes many heart conditions, particularly cardiovascular ischemic events, is thrombosis. The purpose of the current investigation was to determine whether LM leaf extracts had any potential for thrombus breaking. We compared the MSF and its Kupchan fractions at a concentration of 2 mg/mL with the negative control using the positive control’s results. According to our research, test sample thrombolytic activities had positive cascades when compared to positive and negative controls. The results suggested that LM had phytochemicals that are in charge of the activity related to clot lysis.

**Fig. 5.**
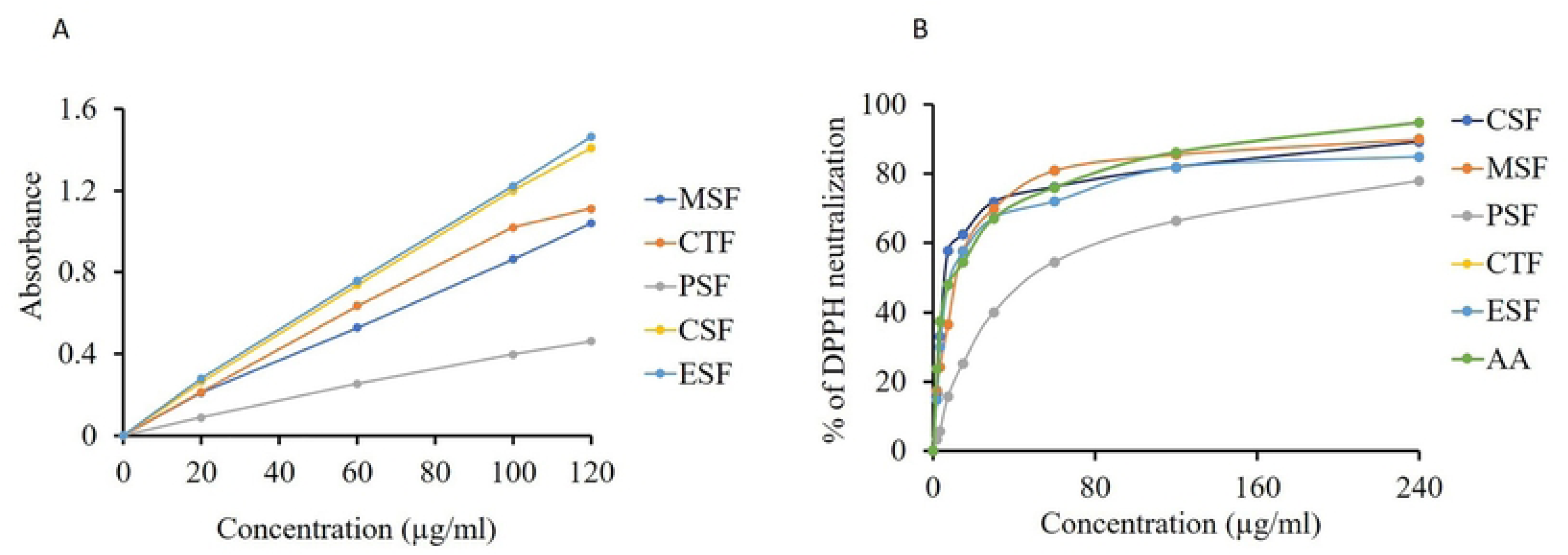
Thrombolytic activities of different fractionates of *LM* leaves.

**Table 7.**
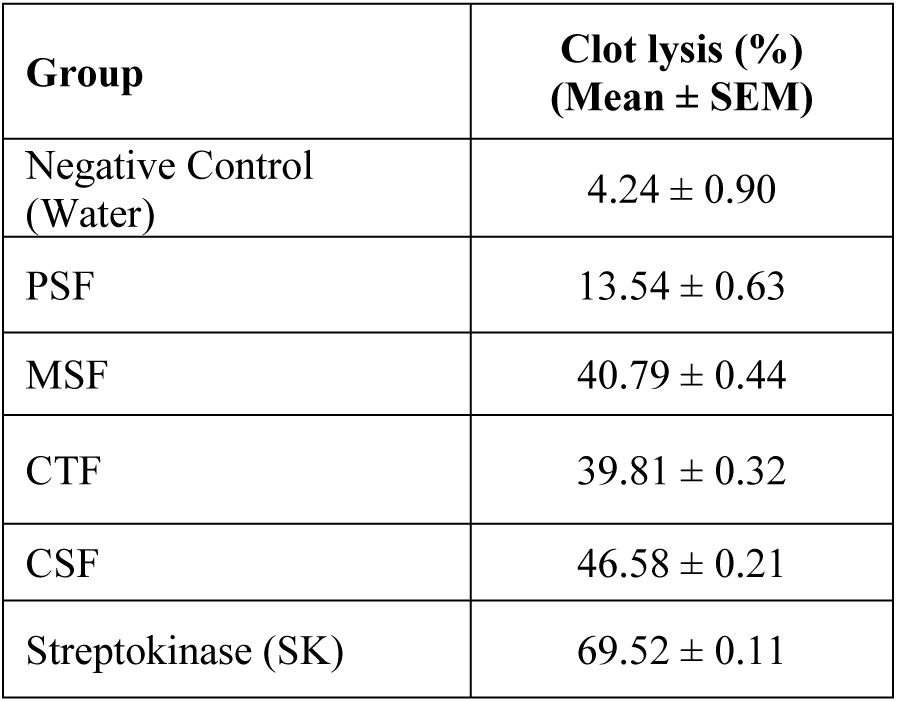
Effect of LM leaves crude extract and Streptokinase on thrombolytic activity after 90 minutes.

| **Group** | **Clot lysis (%)**<br>**(Mean ± SEM)** |
| --- | --- |
| Negative Control<br>(Water) | 4.24 ± 0.90 |
| PSF | 13.54 ± 0.63 |
| MSF | 40.79 ± 0.44 |
| CTF | 39.81 ± 0.32 |
| CSF | 46.58 ± 0.21 |
| Streptokinase (SK) | 69.52 ± 0.11 |

## Conclusion

Our current study’s results allow us to draw the conclusion that the LM leaf extracts shown strong analgesic and anti-diarrheal properties in addition to modest antioxidant thrombolytic activity. The study’s findings indicate that more research may be conducted to identify or separate novel compounds from LM leaf extracts in order to advance the medicinal industry.

## Acknowledgment

The author gratefully acknowledges the Department of Pharmacy, World University of Bangladesh providing full laboratory facilities to conduct this research work and heartfelt gratitude. All the authors have accepted responsibility for the entire content of this submitted manuscript and approved submission.

## Author contributions

Sarder Arifuzzaman and Zubair khalid Labu wrote the manuscript, designed the experiments, and interpreted the data. Samira Karim, and Zubair khalid Labu performed all of the experiments and data collection. Sarder Arifuzzaman performed data analyses. All of the authors were involved in proofreading the manuscript. All of the authors read and approved the final manuscript.

## Declarations

### Availability of data and materials

The datasets generated and/or analysed in this study are available from the corresponding author upon reasonable request.

### Ethics approval and consent to participate

Not applicable.

### Consent for publication

Not applicable.

